# Schisandrin B suppresses colon cancer growth by inducing cell cycle arrest and apoptosis via the CHOP signalling pathway

**DOI:** 10.1101/2023.08.27.554980

**Authors:** Vanessa Anna Co, Hani El-Nezami, Yawen Liu, Bonsra Twum, Priyanka Dey, Paul A Cox, Shalu Joseph, Roland Agbodjan, Mehdi Sabzichi, Roger Draheim, Murphy Lam Yim Wan

**Author notes:** Correspondences Dr Hani El-Nezami School of Biological Sciences, Faculty of Science, Kadoorie Biological Sciences Building, The University of Hong Kong, Pokfulam, Hong Kong, Dr Murphy Lam Yim Wan School of Pharmacy and Biomedical Sciences, Faculty of Science and Health, University of Portsmouth, Portsmouth, United Kingdom.

## Abstract

Colon cancer is among the most lethal and prevalent malignant tumours in the world, and the lack of effective therapies highlights the need for novel therapeutic approaches. Schisandrin B (Sch B), a lignan extracted from the fruit *Schisandra chinensis*, has been reported for its anti-cancer properties. However, no studies to date have been done to characterise the exact molecular mechanisms regarding the anti-tumorigenic effect of Sch B in colon cancer. A comprehensive analysis of the molecular mechanism for the anti-tumorigenic effect of Sch B on human colon cancer cells was performed using combination of Raman spectroscopy, RNA-seq, computational docking and molecular biological experiments. The *in vivo* efficacy was evaluated by a mouse xenograft model. Sch B reduced cell proliferation and triggered apoptosis in human colon cancer cell lines. Raman spectroscopy, computational, RNA-seq, molecular and cellular studies revealed that Sch B activated unfolded protein responses by interacting with CHOP and upregulating CHOP, which thereby induced apoptosis. CHOP knockdown alleviated the Sch B-induced reduction in cell viability and apoptosis. Sch B reduced colon tumour growth *in vivo*. Our findings provide essential background for clinical trials examining the effects of Sch B in patients with colon cancer.

## Introduction

Colorectal cancer (CRC) is the third most common cancer, with over 1.9 million new cases; and the second leading cause of cancer-related death, with over 930,000 deaths [1]. It is estimated that the global burden of CRC will increase to 3.2 million new cases and 1.6 million deaths by 2040 [2]. Several risk factors have been identified over the years, including age, family history, diet, etc. Although CRC is more prevalent in western countries, increasing rates of CRC has been reported in countries where risk was historically low [2]. Current management of CRC is usually chemotherapy accompanied with surgery or radiotherapy [3]. However, these conventional cancer therapies are often accompanied by severe side effects and significant mortality. For example, the response rate of 5-FU in advanced CRC is less than 15%; and long-term administration of 5-FU impairs anti-tumour immune response in patients [4, 5]. Therefore, the identification of a novel but less toxic therapeutic strategy is urgently needed in this patient group.

In recent decades, several lines of scientific evidence suggest that polyphenols that are found abundantly in plant-based foods and drinks, have been shown significant efficacy on cancer development, including chemoprevention and anti-cancer ability. The former is demonstrated in multiple *in vitro*, *in vivo* and epidemiological studies in various cancers, and it is attributed primarily to polyphenols’ potent antioxidative activities [6-8]. The latter is achieved by pro-oxidant actions in cancer cells, which in turn induce cell cycle arrest, apoptosis, and inhibition of cancer cell proliferation [9]. So far, there are a number of studies published exploring the effect of polyphenols on colorectal cell lines or animal models [10-12].

Schisandrin B (Sch B), a lignan extracted from the fruit *Schisandra chinensis*, has been reported for its anti-cancer properties in various cancers, for examples, liver cancer [13], breast cancer [14, 15], ovarian cancer [16], glioma [17], osteosarcoma [18] and gastric cancer [19, 20]. More recently, studies have shown that Sch B was able to prevent or treat colitis-associated colon cancer [21, 22]. Although the antitumor activities of Sch B have been confirmed in various cancer types, the potential mechanism of Sch B seems much more complex, ranging from cell cycle arrest to programmed cell death. For example, in liver cancer, Sch B induced G0/G1 cell cycle arrest and apoptosis by upregulating caspase-3 and Bcl-2 family members in cholangiocarcinoma (CCA), and reduced tumour growth in xenograft mouse model [13]. In breast cancer, Sch B attenuated metastasis in animal model by modulating the epithelial-to-mesenchymal transition (EMT) or STAT3 pathway [14, 15]. In gallbladder cancer, Sch B inhibited cell proliferation and induced apoptosis by regulating apoptosis-related protein expression such as Bax, cleaved capase-3, cleaved caspase-9, Bcl-2 *in vitro* and suppressed tumour growth *in vivo* [23]. In another study by Li *et al*., Sch B is shown to induce glioma cell apoptosis by diminishing mitochondrial membrane potential (ΔΨm). Other studies also reported that Sch B induced apoptosis and cell cycle arrested via Wnt/β□catenin [24], PI3K/AKT [18, 24], NF□κB and p38 MAPK [25] signalling pathways. However, none of these studies have fully characterized the mechanism of the anti-tumorigenic effect of Sch B, especially on colon cancer.

Accordingly, we hypothesized that Sch B would exhibit a significant therapeutic effect against CRC as it does in other cancers. Here we investigated the molecular mechanism for the anti-tumorigenic effect of Sch B on colon cancer through combination of Raman spectroscopy, RNA-seq, computational docking and molecular biological experiments. We propose that Sch B should be further explored as a novel and more specific approach to colon cancer therapy.

## Results

### Sch B inhibited colon cancer growth in human colon cancer cells

The chemical structure of Sch B is depicted in Fig. 1a. The CCK-8 assay was performed to determine the effects and concentrations of Sch B on multiple CRC cell lines of different stages. Sch B displayed a concentration-dependent effect on all the CRC cell lines (*P* < 0.05), except for Caco-2. Although Sch B also lowered the cell viability of the normal cell line CCD841 CoN, the change was not statistically significant. Based on the half maximal inhibitory concentration (IC_50_) calculated from the CCK-8 assay, HCT116, HT29 and SW620 cell lines were found to be the most sensitive to Sch B treatment and were therefore used for subsequent experiments (Fig. 1b-c, Supplementary Fig. 1). Trypan blue exclusion and BrdU-ELISA assays were also performed to support the results from CCK assay (Fig. 1d, f). In addition, the long-term anti-proliferative effect of Sch B on CRC cells were assessed by clonogenic survival assay. Sch B reduced the colony-forming ability significantly of HCT116, HT29 and SW620 after 48-hour treatment. The loss of cell viability was irreversible, as shown after 14 days (Fig. 1e).

**Fig. 1.**
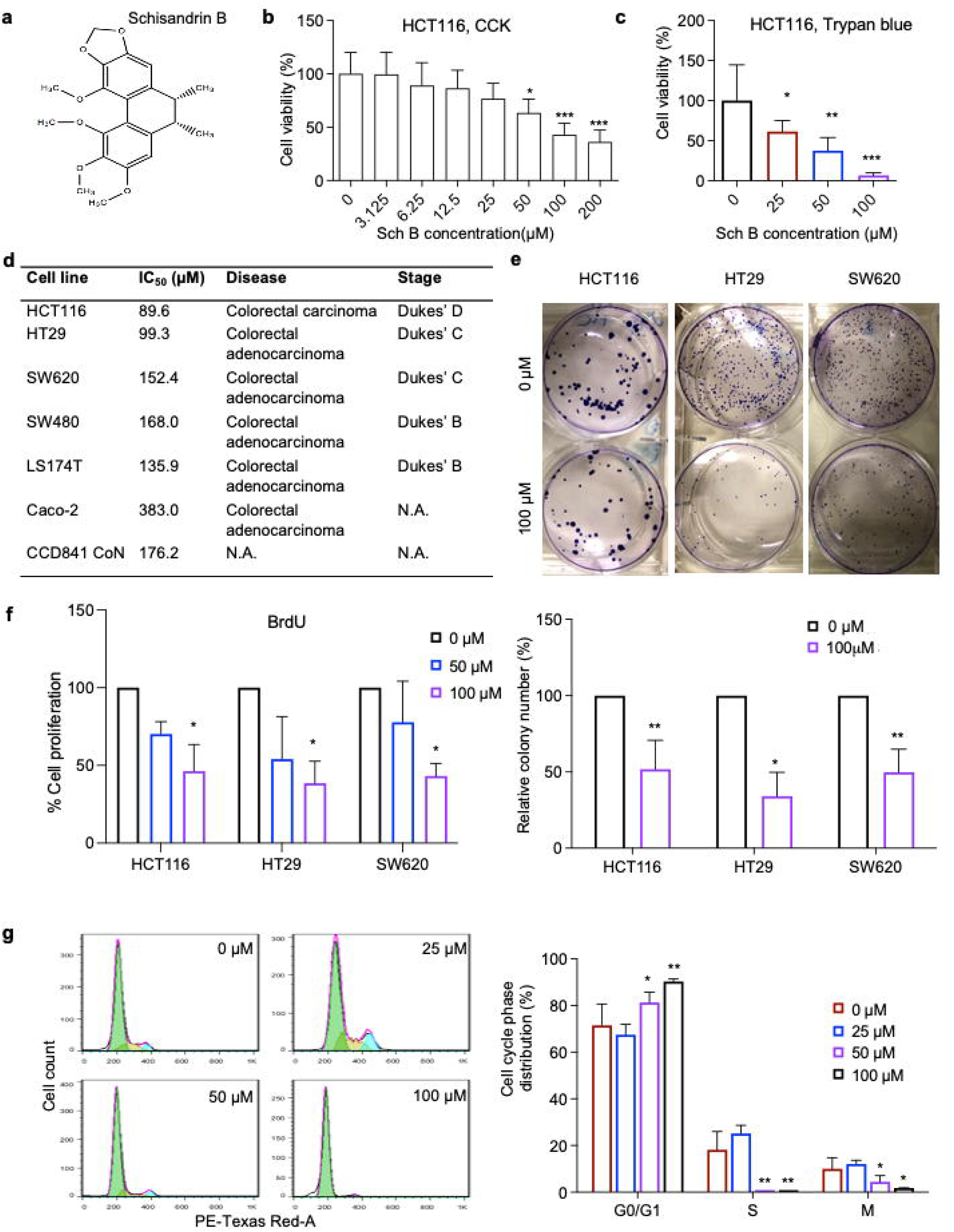
Schisandrin B inhibited the proliferation of human colon cancer cells. **a,** The chemical structure of Schisandrin B (Sch B). **b-c**, The viability of human HCT-116 colon cancer cells incubated with various concentrations of Sch B for 48 hours, quantified by the Cell Counting Kit-8 (CCK-8) assay (**b**) and trypan blue exclusion assay (**c**) (*n* = 3 experiments). **d,** Half maximal inhibitory concentration (IC_50_) of human colon cancer cell lines HCT116, HT-29, SW480, SW620, Caco-2 and LS174-T, and normal human colon cell line CCD 841 CoN. **e,** Colony assay showing long-term effects of Sch B (100 μM) on the colony formation potential of HCT116, HT29 and SW620 cells (*n* = 4-5 experiments). **f**, Evaluation of BrdU incorporation as an index of DNA synthesis, after Sch B (50 and 100 μM) treatment (*n* = 3 experiments). **g,** Cell cycle analysis of HCT116 cells treated with 0, 25, 50 and 100 μM Sch B for 48 hours (*n* = 6 experiments). Values were presented as mean ± SD, analysed by Kruskal–Wallis test with Dunn’s correction (**b, f**), One-way ANOVA with Holm-Šídák’s multiple comparisons (**c, g**) or Mann–Whitney U-test (**e**). \**P* < 0.05, \*\**P* < 0.01, \*\*\**P* < 0.001, compared with the control (i.e., 0 μM Sch B).

We further analysed the cell cycle distribution of the Sch B-treated HCT116 cells. Results showed that Sch B treatment led to a decrease in cell proportion at S and M phases and an accumulation in cell proportion at G0/G1 phase (Fig. 1g). These data indicate that Sch B arrests HCT116 cells at the G0/G1 phase of the cell cycle and subsequently blocks cell growth.

### Sch B promoted apoptosis in human colon cancer cells

Cell apoptosis was analysed by using Annexin V/PI staining. Results from flow cytometry showed an accumulation of apoptotic and necrotic cells in the human colon cancer cells exposed to increasing concentrations of Sch B (Fig. 2a).

**Fig. 2.**
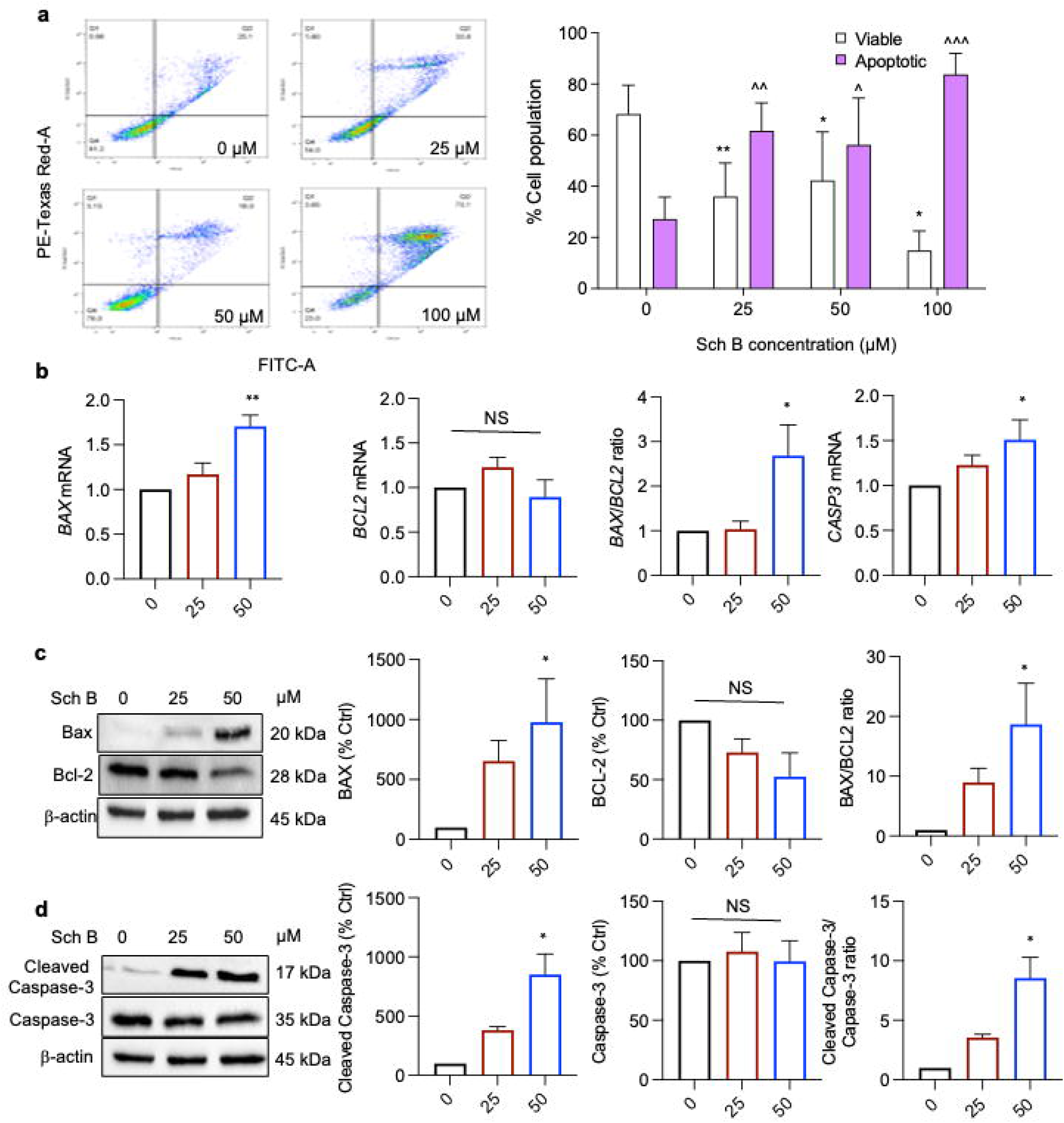
Sch B promoted apoptosis in HCT116 cells. **a,** Evaluation of apoptosis rates of HCT116 cells incubated with 0, 25, 50 and 100 μM Sch B for 48 hours by Annexin V/PI flow cytometry (*n* = 5 experiments). **b,** mRNA expression levels of *BAX*, *BCL2*, *CASP3* in cells treated with 0, 25 and 50 μM Sch B for 48 hours. The mRNA expression levels were normalised to GAPDH. **c-d,** Immunoblot analysis of BAX, BCL-2, cleaved CASP3 and CASP3. The protein levels were normalised to β-actin (*n* = 3 experiments). Values were presented as mean ± SD, analysed by One-way ANOVA with Holm-Šídák’s multiple comparisons (**a**) or Kruskal–Wallis test with Dunn’s correction (**b-d**). *, ^*P* < 0.05, **, ^^*P* < 0.01, ***, ^^^*P* < 0.001, compared with the control (i.e., 0 μM Sch B). NS = no significance.

qPCR and Western blot were also used to analyse the mRNA and protein levels of several important apoptosis-related markers (*BAX*, *BCL2* and *CASP3*) (Fig. 2b-d). Results of qPCR showed that the mRNA levels of apoptotic genes *BAX* and *CASP3* were elevated in the Sch B-treated cells but no change in anti-apoptotic gene *BCL2* mRNA was observed. Since the balance between pro-apoptotic proteins and anti-apoptotic proteins of Bcl family plays a key role in regulation of intrinsic pathway of cell apoptosis, we calculated the BAX/BCL2 ratio as a proper indicative of cell apoptosis. Sch B significantly increased BAX/BCL2 ratio in a concentration-dependent manner (Fig. 2b). Similarly, Sch B treatment significantly increased BAX and cleaved (active) caspase-3 protein level while there was no difference in pro-caspase-3 protein levels among all treatment groups. The increases in BAX/BCL2 and active caspase-3/pro-caspase-3 ratios indicated that Sch B might induce human colon cancer cell death via promoting apoptosis (Fig. 2c-d).

### Raman spectroscopy revealed changes in molecular signatures in cells after Sch B treatment

To uncover the underlying mechanism for the anti-cancer effects of Sch B, we first explored the uptake of Sch B and effect on human colon cancer cells by using non-destructive, label-free Raman spectroscopy. The Raman spectrum of Sch B powder sample was analysed by the commercial Renishaw Raman spectrometer with several signature peaks occurred at 680, 714 and 1420 cm^-1^ (depicted with pink bars) (Supplementary Fig. 2a). Cell lysates and cell-free supernatants were obtained from human colon cancer cells exposed to 50 µM Sch B (i.e., 20 µg/mL) and analysed by an in-house compact Raman spectrometer. The spectra of cell lysates and cell-free supernatants from cells cultured in the presence or absence of Sch B is shown in Fig. 3b. A slight Raman shift was detected in cell lysates and cell-free supernatants containing Sch B compared to Sch B powder, where the signature Raman peaks occurred at 683, 719 and 1420 cm^-1^ (depicted with pink bars). Such shift in peak positions may be due to the differences in the spectral resolution of the two Raman systems and/or solvation caused by Sch B in cell media. However, these Raman peak positions were absent in the cell lysate from cells without Sch B treatment, confirming that these peaks are related to Sch B.

**Fig. 3.**
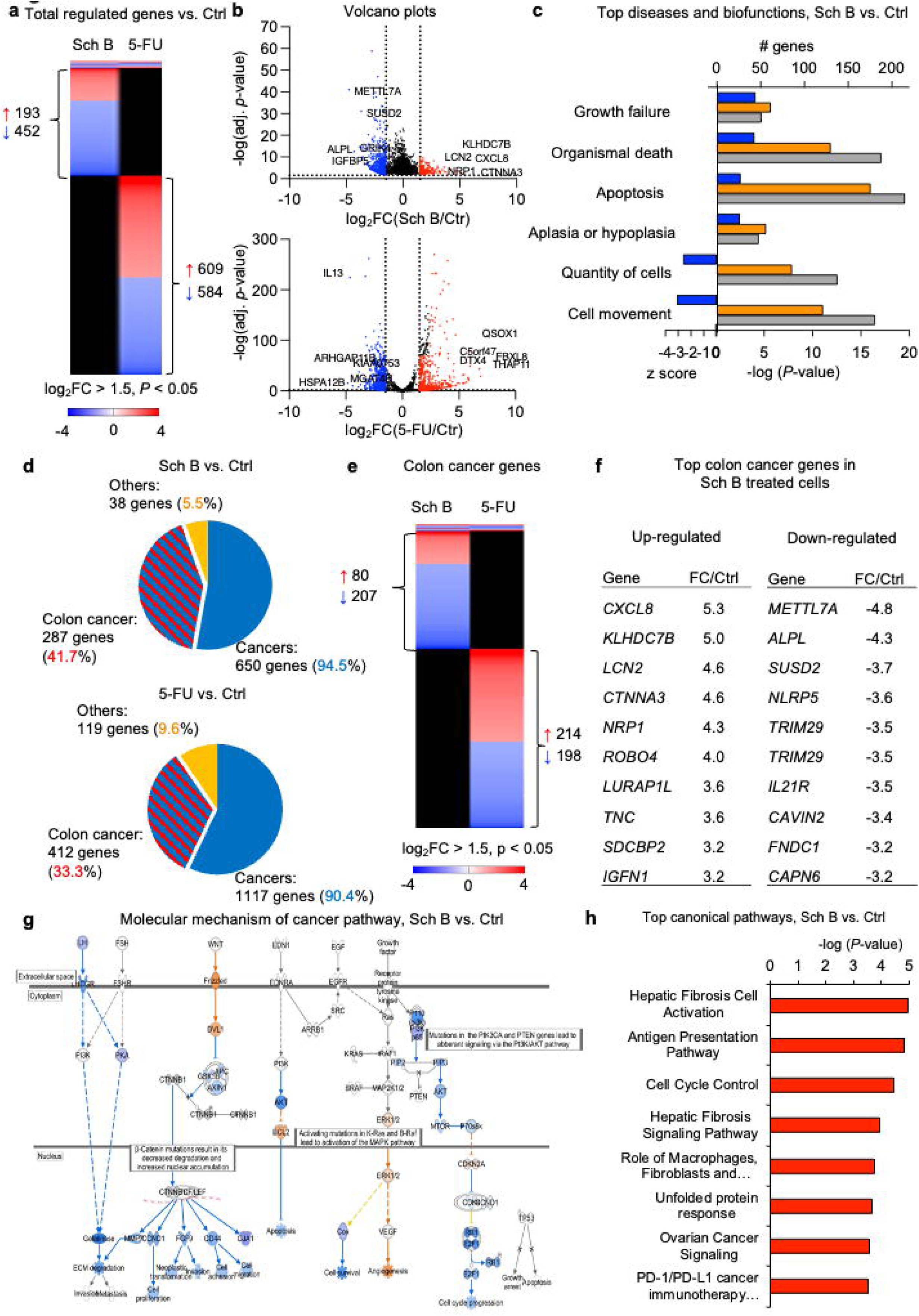
RNA sequencing analysis of HCT116 cells treated with Sch B or 5-FU. **a,** Heatmap comparing gene expression profiles of cells after exposure to 50 µM Sch B or 100 µM 5-FU compared to untreated control. (red, upregulated; blue, downregulated; black, not regulated; cutoff log_2_FC□>□1.5, *P* < 0.05, *n*□=□1 experiment). **b,** Volcano plots showing differentially expressed genes in Sch B or 5-FU treated cells. **c,** Biofunction analysis showed Sch B affected genes related to cell growth, cell movement, organism death and apoptosis. Blue, Z-score; orange, *P*-value; grey, no. of genes. **d,** Pie chart of genes regulated in response to Sch B or 5-FU treatment. Gene categories were identified by biofunction analysis. **e,** Heatmap comparing colon cancer related gene expression profiles of cells after exposure to 50 µM Sch B or 100 µM 5-FU compared to untreated control. **f,** Top colon cancer related genes in cells treated with Sch B. **g,** IPA analysis showing inhibition of the cancer pathway in Sch B treated cells. Orange, up-regulated genes; blue, down-regulated genes. **h,** Top canonical pathway affected by Sch B.

A visual comparison of spectral intensities further showed differences in the Raman spectra of the cell lysates at 0- and 48-hour time points (Supplementary Fig. 2b). The peaks pertaining to Sch B and cellular components are depicted with pink and red bars, respectively. It is interesting to note that Raman peaks were observed at 882 and 1457 cm^-1^ in the spectrum of cell lysates obtained at 0 hour but not 48 hours, which are often associated with C-C skeletal vibrations and CH_2_ scissoring and CH_3_ deformation vibrations, suggesting presence of aliphatic amino acids and/or glucose, respectively. The identified peaks may be considered as potential indicators of the drug effect on the metabolism or death of the cells, and further experiments are required to verify this.

Furthermore, cellular uptake is another important contributor to the efficiency and biological activity of novel drugs. To monitor the Sch B uptake to the cells, relative intensities of the Raman peaks of Sch B (683, 719 and 1420 cm^-1^) were measured after 0 and 48 hours of Sch B treatment. At the 0-hour time point, a rapid uptake (17%) of Sch B into the cells occurred. At 48 hours, the level of Sch B uptaken by the cells were significantly increased to about 42%, which corresponds to approximately 12.6 µg of Sch B taken up by the cells (initial cell number 2.16×10^6^, 5.83 pg Sch B per cell).

### Sch B induced differential gene expression in cancer cells

In order to identify the genome-wide alterations in the transcriptome of human colon cancer cells treated with Sch B, total RNA was extracted from cells and was subjected to RNA-seq. Gene expression profiles were compared between untreated cells and cells after exposure to Sch B. 5-FU-treated cells served as positive controls (Fig. 3).

Our RNA-seq results revealed that major differences between Sch B treated and 5-FU treated cells compared to untreated cells. There were 193 upregulated genes and 452 downregulated genes for the Sch B group (cutoff log_2_FC > 1.5, *P* < 0.05) (Fig. 3a). The top regulated genes included *METTL7A*, *ALPL*, *SUSD2*, *IGFBP5*, *CORO1A*, *CXCL8*, *LCN2*, *NRP1* and *CTNNA2* that are involved in tumour proliferation, angiogenesis, migration, metastasis and tumour immunity. In 5-FU treated cells, 609 genes were upregulated, and 584 genes were downregulated. The top regulated genes included *KIAA0753, IL13, HSPA12B, MGAT4B, THAP11, QSOX1, DTX4* and *FBXL8* which are related to cell movement, cancer invasion and metastasis, drug resistance and inflammation and immune-overactivation (Fig. 3b). Functional analysis revealed that Sch B affected genes related to cell growth, cell movement, organism death and apoptosis, which were consistent with the results from previous cell death and functional assays (Fig. 3c).

While only 43 genes were commonly regulated between Sch B and 5-FU treated cells which indicated that Sch B and 5-FU had very different targets for gene regulation, Sch B treatment was also as efficient as 5-FU as defined by strong effects on cancer gene networks (Fig. 3d-f). 95% of all regulated genes in RNA extracts from cells treated with Sch B were cancer-related. This included a subset of colon cancer-related genes, which were inhibited (41.7%) (Fig. 3d-f).

The molecular mechanisms of cancer pathway genes were deactivated, including *COX, GJA1, CD44, FGF9* and *CCND1* (Fig. 3g). Further pathway analysis of the RNA-seq data also detected several major pathways regulated by Sch B, which included hepatic fibrous and ovarian cancer signalling, cell cycle control, unfolded protein response and PD-1/PD-L1 cancer immunotherapy (Fig. 3h).

### Sch B promoted ER stress-dependent apoptosis in colon cancer cells via activation of CHOP signalling

Unfolded protein response (UPR) is a known cellular defence mechanism in response to endoplasmic reticulum (ER) stress and is implicated in cancer progression and pathogenesis. Our results showed unfolded protein responses to be one of the top canonical pathways regulated by Sch B. Analysis of genes involved in the unfolded protein response pathway revealed DDIT3 as the top regulated gene that was up-regulated by Sch B treatment (Fig. 4a).

**Fig. 4.**
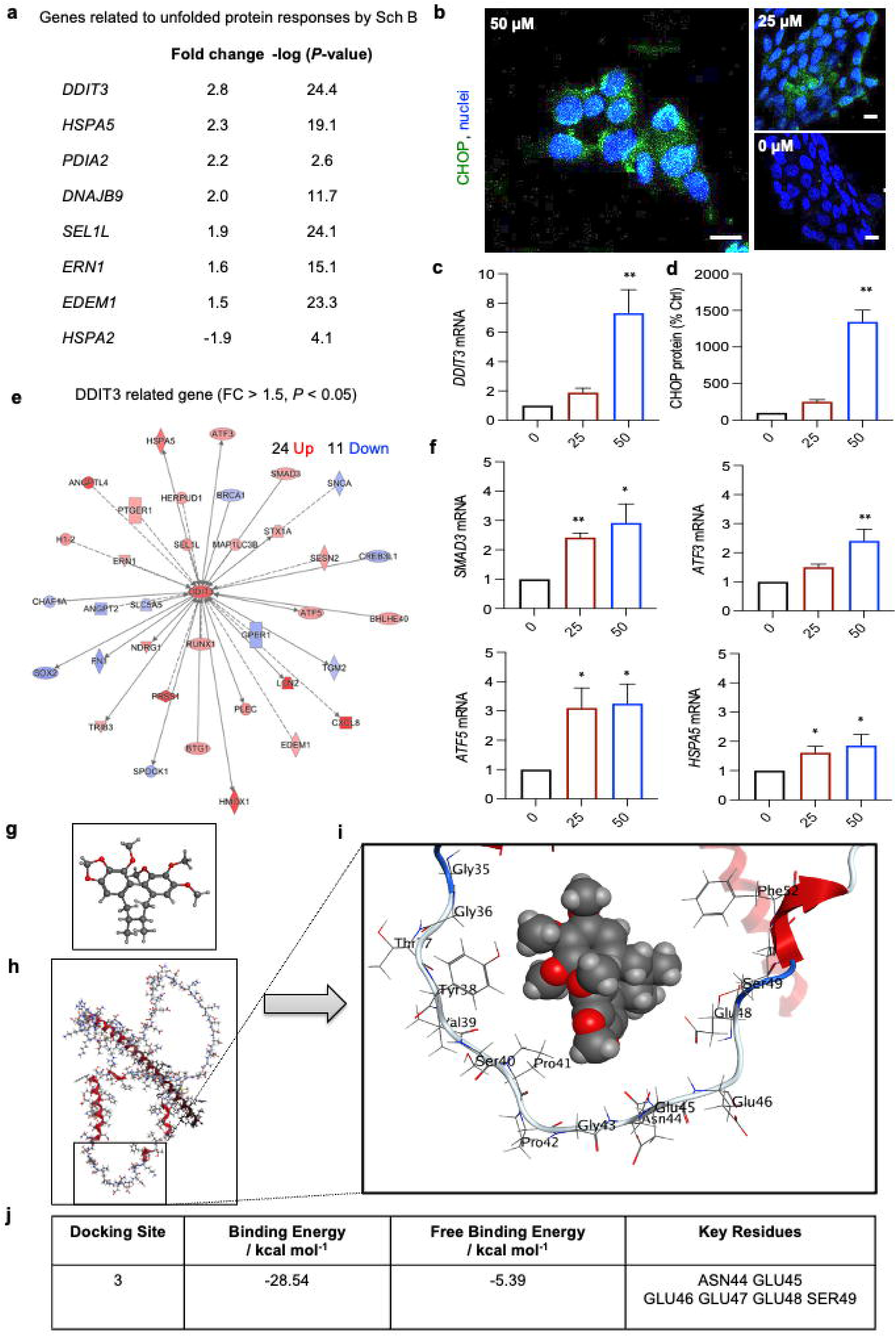
Sch B activated CHOP signalling pathway of the unfolded protein response. **a,** Ingenuity Pathway Analysis (IPA) analysis identified *DDIT3* as one of the top upregulated genes. **b,** Confocal imaging of CHOP protein (green, CHOP; blue, nuclei; scale bars, 10 μm) (*n* = 3 experiments). **c,** mRNA level of DDIT3 by qPCR (*n* = 4 experiments). **d,** Protein expression level of CHOP by immunofluorescence staining (right). For confocal image quantification, 50□cells were counted per sample (*n* = 3 experiments). **e,** *DDIT3* and DDIT-3 dependent gene network. Red, up-regulated genes; blue, down-regulated genes; cutoff log_2_FC□>□1.5, *P* < 0.05. **f,** mRNA expression of DDIT3 dependent gene expression quantified by qPCR (*n* = 4 experiments). Values were presented as mean ± SD, analysed by Kruskal–Wallis test with Dunn’s correction (**c, d, f**). \**P* < 0.05, \*\**P* < 0.01, \*\*\**P* < 0.001, compared with the control (i.e., 0 μM Sch B). **g-j,** Proposed molecular model of Sch B binding with CHOP. **g,** The 3D chemical structure of Sch B. **h,** The CHOP protein structure. **i**, The energy minimised location for the Sch B molecule docked with the CHOP protein. The rectangle in (**h**) shows the part of the CHOP structure represented in (**i**). **j,** The docking site with the lowest binding energy and key residues involved in the interaction.

To confirm the results of RNA-seq, we performed real-time qPCR and immunofluorescence staining of DDIT3 (CHOP) expression (Fig. 4b-c). Consistent with RNA-seq data, both mRNA and protein level of DDIT3 (CHOP) was remarkably up-regulated in cells following Sch B treatment. Further transcriptomic analysis identified a network of DDIT3 (CHOP)-dependent gene being up-regulated (Fig. 4e). qPCR analysis confirmed that mRNA expression of several CHOP-dependent genes, including *SMAD3*, *ATF3*, *ATF5* and *HSPA5* that were significantly up-regulated in Sch B treated cells (Fig. 4e).

To investigate the interaction between the CHOP protein and the Sch B molecule, molecular docking was performed (Fig. 4g-j). Nine potential binding sites were identified using the Site Finder tool within the Molecular Operating Environment (MOE) program [26]. Of these sites, two had significantly higher binding energies than the other seven (Supplementary Table 2). These two sites (sites 3 and 7) are very close together in the structure since they share common residues (GLU46, GLU47 and SER49). The energy minimised location for the Sch B molecule at site 3 is shown in Fig. 4i-j.

Furthermore, it is well documented that activation of CHOP during ER stress can induce cell cycle arrest and oxidative stress, leading to cell death and apoptosis [27]. To determine the role of ER stress in the Sch B-induced growth inhibition, we performed siRNA-mediated silencing of CHOP in HCT116 cells (Fig. 5a-b). Using CCK-8 assay and Annexin V/PI staining, we found that CHOP knockdown moderately restored the Sch B-induced changes in cell viability and Sch B-induced apoptosis was attenuated (Fig. 5c-d). These results suggest that CHOP activation involves in the regulation of Sch B-triggered responses (Fig. 5e).

**Fig. 5.**
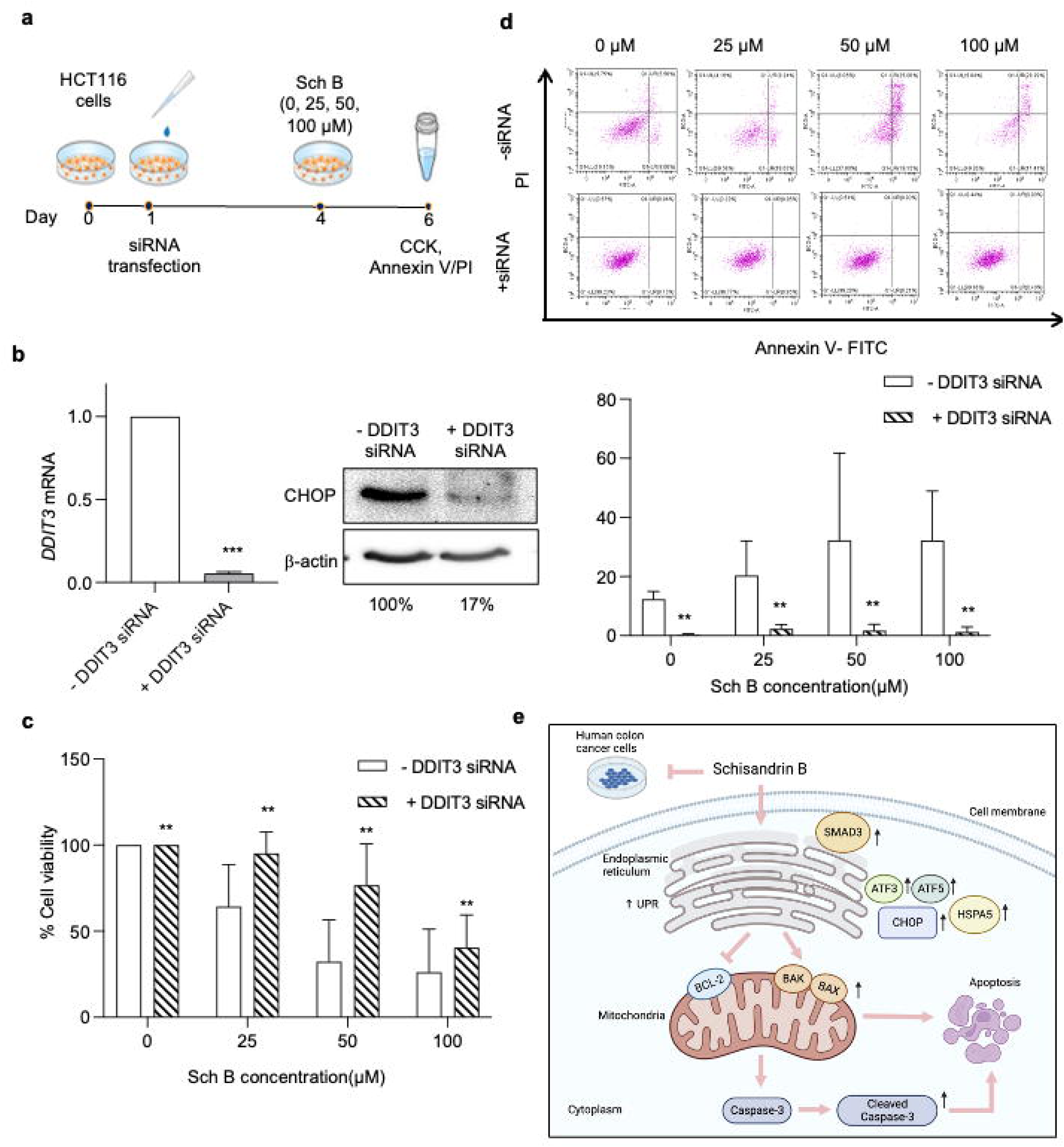
Sch B promoted CHOP-dependent apoptosis in human colon cancer cells. **a,** Schematic of the protocol used to treat HCT116. Cells were treated with siRNA for 72 hours and then treated without or with 0, 25 and 50 µM Sch B for 48 hours. Cells were harvested for CCK assay and Annexin V/PI flow cytometry. **b,** mRNA and protein expression levels of DDIT3/CHOP in siCHOP-transfected cells. The mRNA and protein expression levels were normalised to GAPDH and β-actin, respectively (*n* = 2 experiments). The mRNA and protein levels were normalised to GAPDH and β-actin, respectively. **c,** The viability of siCHOP-transfected cells incubated with 0, 25, 50 and 100 μM Sch B for 48 hours, as quantified by CCK assay (*n* = 3 experiments). **d,** Evaluation of apoptosis rates of siCHOP-transfected HCT116 cells by Annexin V/PI flow cytometry (*n* = 3 experiments). **e,** Schematic of CHOP-dependent induction of apoptosis by Sch B. Values were presented as mean ± SD, analysed by Wilcoxon matched-pairs signed-rank test (**b**), One-way ANOVA with Holm-Šídák’s multiple comparisons (**c, d**). \**P* < 0.05, \*\**P* < 0.01, \*\*\**P* < 0.001, compared with the control (i.e., 0 μM Sch B).

### Sch B suppressed colon cancer growth in vivo

To characterize the *in vivo* anti-tumour effect of Sch B, we generated nude mice bearing HCT116 cell xenografts. Mice were treated perorally every other day with Sch B at dosages of 50 mg/kg body weight for one week. 5-Fluorouracil (5-FU)-injected mice served as positive control (Fig. 6a). Results showed that tumour volume was significantly reduced in Sch B-treated mice compared to the sham-treated mice. In addition, tumour weight showed a reduction in the Sch B-treated mice, while the body weight remains unchanged between the Sch B- and sham-treated groups (Fig. 6b-d). Since diarrhoea is one of the side-effects reported in 5-FU treatment, the animals were monitored daily for diarrhoea since the start of the treatment. Sham- and Sch B-treated mice did not report any incidence of diarrhoea. The animals in the 5-FU group had a significantly higher cumulative diarrhoea assessment score compared to the other groups since day four (Fig. 6e). Histopathological examinations were further performed by staining with haematoxylin-eosin (H&E) to examine any morphologic changes relevant to apoptosis/necrosis in tumour tissues. The results indicated a lower tumour cell density in the Sch B and 5-FU groups compared to the sham group. Tumour necrosis was found in both Sch B- and 5-FU-treated tumour tissues and inflammatory cell infiltration was also observed in Sch B-treated tumour tissues (Fig. 6f).

**Fig. 6.**
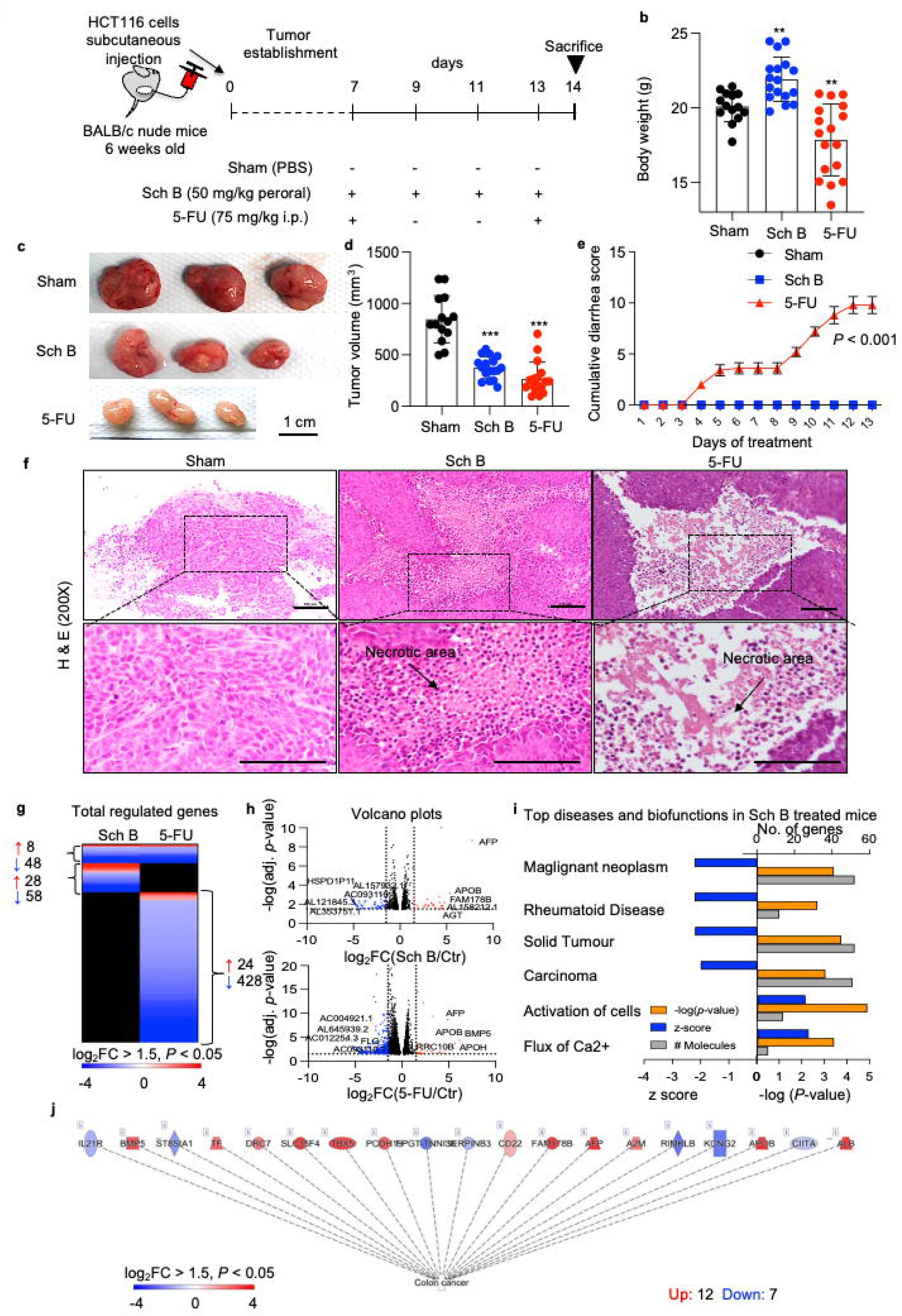
Therapeutic efficacy of Sch B in a mouse xenograft model of human colon cancer. **a,** Schematic of the protocol used to treat HCT116 colon cancer xenografts in nude mice. Six-week-old male BALB/C nude mice injected with HCT116 cells subcutaneously. One week after tumour injection, the mice were given either perorally with Sch B (50 mg/kg) every other day or injected intraperitoneally once a week with 5-FU (75 mg/kg). The treatment lasted for 14 days. **b,** Body weight of mice in different treatment groups. **c,** Representative images of tumour from each treatment group. Scale bars, 1 cm. **d,** Tumour volume. **e,** Mice was monitored daily after treatment to observe for any diarrhoea. A diarrhoea assessment score was given to each mouse based on wetness and hardness of the stool. 0 = normal (normal stool or absent); 1 = slight (slightly wet and soft stool); 2 = moderate (wet and unformed stool with moderate perianal staining of the coat); 3 = severe (watery stool with severe perianal staining of the coat). **f,** Microscopic examination of hematoxylin and eosin (H&E) stained tumour sections (200x). Scale bars, 100 µm. The arrows indicated the areas of necrosis. g, Heatmap comparing gene expression profiles of tumours from Sch B and 5-FU treated mice compared to untreated control. (red, upregulated; blue, downregulated; black, not regulated; cutoff log_2_FC□>□1.5, *P* < 0.05, *n*□=□3 mice per group). **h,** Volcano plots showing differentially expressed genes in tumours from Sch B or 5-FU treated mice. **i,** Top diseases and biofunctions of significantly regulated genes by Sch B. Blue, Z-score; orange, *P*-value; grey, no. of genes. **j,** Several colon cancer gene were significantly regulated in tumours obtained from mice receiving Sch B compared to untreated control.

To characterize the tumour response to Sch B, total RNA derived from tumour tissues was subjected to RNA-seq. Gene expression profiles were compared between the sham treated group and mice receiving Sch B. Tumour tissues from 5-FU-treated mice served as positive controls (Fig. 6g-j).

Similar to our *in vitro* studies, RNAseq results revealed a profound difference in gene expression profiles between Sch B and 5-FU treatment groups compared to sham. There were 36 up-regulated genes and 106 down-regulated genes for the Sch B group (cut-off log_2_FC 1.5, *P* < 0.05) (Fig. 6g). The top regulated genes included long non-coding RNAs (lncRNAs; *AL157932.1, AC093110.1, AL353751.1*), *HSPD1P11, AFP, APOB, FAM178B* and *AGT*. In 5-FU treatment group, 32 genes were upregulated, and 476 genes were downregulated. The top regulated genes included *BMP5, APOB, AFP, APOH, LRRC10B, FLG* as well as some lncRNAs (*AC012254.3, AL645939.2, AC004921.1*) (Fig. 6h). Despite a relatively low number of genes regulated by Sch B, cancer biofunctions were markedly attenuated (Z-score > -2) in Sch B-treated mice compared to the sham group (Fig. 6i). A subset of colon cancer genes was specifically regulated (Fig. 6j).

Further pathway analysis revealed several major pathways regulated by Sch B, including LXR/RXR activation, acute phase response signalling, clathrin-mediated endocytosis signalling and production of NOs and ROS in macrophages. However, unfolded protein response signalling was not significantly affected in Sch B-treated tumour tissues (Supplementary Fig. 3).

## Discussion

Conventional cancer therapies are often accompanied with various side effects and organ toxicities, resulting in a lower quality of life for patients [28]. While chemotherapy remains the mainstay of cancer treatment, it often incur high costs [29]. It is estimated the monthly cost of 5-FU-based chemotherapy to be USD2370 to USD4665 for each patient, depending on the administration method [30]. As such, there is an urgent need to develop less costly alternatives to chemotherapy for CRC that could lead to improved clinical outcomes.

Our data have discovered an entirely new approach which offers a safe and cost-effective alternative to human colon cancer treatment. Sch B, a lignan found in the fruit of *Schisandra chinensis* (also known as Five-flavour berry), offers astonishing properties, and goes beyond current cancer-specific therapies in the following aspects: 1) It is a natural polyphenol with high tumour-killing capacity and high degree of specificity; 2) It treats different stages of colon cancers, especially more effective for the late stage of colon cancers; 3) Sch B has shown very low toxicity against normal cells compared to current available drugs.

Furthermore, this study was the first to provide a comprehensive understanding of the molecular mechanism for the anti-tumorigenic effect of Sch B on colon cancer through combination of Raman spectroscopy, RNA-seq, computational docking and molecular biological experiments. Mechanistic investigation reveals Sch B treatment triggered apoptosis in the colon cancer cells through activation of ER stress pathway. Our study has shown that Sch B upregulated UPR markers, such as DDIT3 and HSAP78, etc., indicating that that Sch B induced ER stress in CRC cells. ER stress is regularly reported in cancer cells [31]. In response to ER stress, URP occurs to restore homeostasis of the ER folding environment. It can either promote cell survival or cell death in tumour cells, depending on the cell status [32]. In cells with prolonged ER stress, C/EBP homologous protein (CHOP, also known as DDIT3) is upregulated and subsequently induces apoptosis in cancer cells via multiple signalling pathways [33]. Previous studies have demonstrated the pro-apoptotic and anti-proliferative effects of activated CHOP signalling in cancer cells [34, 35]. In mitochondria-dependent apoptosis, CHOP regulates the expression of BCL-2 family proteins and TRB3 protein [36]. Overexpression of CHOP can induce translocation of BAX protein from the cytosol to the mitochondria, resulting in cell death [37]. Consistent to our findings, we demonstrate that Sch B could effectively bind to the active sites of CHOP. It is interesting to note that the position of Sch B is away from the two functional domains in the CHOP protein, the N-terminal transcriptional activation domain and the C-terminal basic-leucine zipper domain. This would seem to be consistent with the Sch B molecule potentially activating the protein rather than inhibiting it. Furthermore, we showed that in cells where CHOP is silenced, the efficacy of Sch B against CRC cells was significantly reduced, compared to non-transfected cells. These data suggest that Sch B may induce cell death through activation of CHOP.

Therapeutic efficacy of Sch B on colon cancer was also confirmed in mouse xenograft model in this study. While the unfolded protein response was not significantly regulated by the Sch B in mouse model, it could be due to differences in dosage and administration method, as well as the larger variation among animal-derived samples. In addition, the discovery of the role of unfolded protein response in cancer progression is relatively new and its mechanisms are still not fully understood [38]. Studies have reported both pro-carcinogenic and anti-carcinogenic effects of ER stress, but it is unclear what factor determines the role and the outcome of ER stress in cancer [39, 40]. Moreover, one proposed CHOP-induced apoptosis was via upregulating ERO1α, which can increase ROS production, and trigger the release of Ca2+ from the ER to the mitochondria, thereby causing oxidative stress and cell apoptosis [41]. The increased concentration of Ca2+ in the cytosol then activates CaMKII and NOX2, and promotes ROS production, creating a positive feedback pathway [35]. An upregulation of flux of Ca2+ was observed in transplanted tumour cells from the mouse model. Also, there is substantial evidence that the activation of CHOP is linked to favourable outcome of cancer treatment. There are multiple pathways in which CHOP induces apoptosis. So far, the mitochondria-dependent pathway, PERK-ATF4-CHOP pathway and CHOP-induced ROS have been identified [35]. It is possible that Sch B-induced CHOP inhibit tumorigenesis via different CHOP-dependent pathways in the *in vitro* and *in vivo* models. Therefore, it might be worth investigating interaction of Sch B and CHOP signalling by looking at downstream regulators of CHOP signalling in the CRC cells and transplanted tumour tissues, respectively.

Overall, we demonstrated that Sch B induced CHOP-dependent apoptosis and inhibited cell proliferation and tumour growth *in vitro* and *in vivo.* The promising anti-cancer results indicate a potential and effective therapeutic strategy that uses Sch B for CRC treatment. The combination of Raman spectroscopy, RNA-seq, computational docking and molecular biological experiments provide comprehensive information about the mechanism for the anti-colon cancer effects of Sch B, making it possible to identify and develop novel molecular and biological targets.

## Materials and Methods

### Chemicals and reagents

Dulbecco’s Modified Eagle’s Medium (DMEM), Foetal Bovine Serum (FBS), Trypsin-EDTA and other cell culture reagents were purchased from Gibco-Life Technology (Eggenstein, Germany). Sch B was purchased from MedChem Express (Monmouth Junction, NJ, USA), and 5-fluorouracil (5-FU) and crystal violet from Sigma-Aldrich (St. Louis, MO, USA). They were dissolved in dimethylsulfoxide (DMSO; Sigma) and stored at -20°C before use. In addition, phosphate buffered saline (PBS), sodium biocarbonate, diethylpyrocarbonate (DEPC) and chloroform were purchased from Sigma. Ammonium persulfate, N, N, N’, N’-tetramethyl-ethane-1,2-diamine (TEMED), acrylamide, resolving gel buffer, stacking gel buffer, 10% (w/v) Tween 20 and 20% (v/v) sodium dodecyl sulfate (SDS) were purchased from Biorad (Richmond, CA, USA). Absolute ethanol, and isopropanol were obtained from Merck (Darmstadt, Germany). RNAisoPlus, PrimeScript RT reagent kit with gDNA Eraser and TB Green® Premix Ex Taq^TM^ were purchased from Takara (Otsu, Japan). Molecular Probes™ Dead Cell Apoptosis Kits with Annexin V for Flow Cytometry, Propidium iodide, GeneJET RNA Purification kit, Histomount were from Thermo Fisher Scientific (Dreieich, Germany). HiScript^TM^ RT SuperMix for qPCR and AceQ qPCR SYBR Green Master Mix were obtained from Vazyme Biotech Co. (Piscataway, NJ, USA).

### Cell culture

Human colorectal adenocarcinoma HCT-116, HT-29, SW480, SW620, Caco-2 and LS174-T, and normal human colon cells CCD 841 CoN were cultured in Dulbecco′s modified eagle′s medium (DMEM) supplemented with 10% fetal bovine serum (FBS) (v/v) and maintained at 37°C under a humidified atmosphere, with 5% CO_2_. Cells within 10 passages were used for all assays. All cells were screened for mycoplasma contamination with a MycoAlert mycoplasma detection kit (Lonza, Basel, Switzerland) prior to use.

### Cell viability assay (CCK-8 assay)

Cell Counting Kit-8 (CCK-8) assay (Dojindo, Kunamoto, Japan) was used to assess cell viability after exposure to Sch B for the indicated time points. 10^4^ cells/well were seeded into 96-well culture plates (Corning, NY, USA) and allowed to adhere for 24 hours. Then, cells were then treated with different concentrations of Sch B (0-200 µM) for 48 hours. 10µL of CCK-8 was added to the wells after treatment and incubated at 37°C for one hour. Absorbance of the wells was measured at 450nm. The viability of human colon cancer cells after treatment was expressed as the proportion of optical density (OD) compared to control (untreated cells). Based on the cell viability assay results, HCT-116 cells were found to be the most sensitive to Sch B treatment and were therefore selected to proceed in other assays.

### Cell cycle flow cytometry

Cells were seeded to 60mm culture dishes and treated with Sch B for 48 hours. Cells washed with PBS and harvested by using Trypsin-EDTA (0.05%). DMEM containing 10% FBS was used to inactivate the trypsin. The cells were centrifuged at 1000 rpm for five minutes to remove the supernatant and were washed with PBS twice. Then, the cells were fixed in ice-cold 70% ethanol and stored at -20°C. Fixed cells were spun down at 500 x g for five minutes at room temperature and washed with PBS. The supernatant was aspirated, and cells were stained with 500µL PI staining solution for 30 minutes. BD Bioscience FacsAria III flow cytometer (San Jose, CA, USA) was used to analyse the fluorescent output on the FITC and PI emission wavelengths. FlowJo 7.6 was used for data analysis.

### Annexin V/PI flow cytometry

Cells with or without transfection were seeded to 60mm culture dishes or 96-well plates and treated with Sch B or 5-FU for 48 hours. The spent medium was collected, and the cells were harvested using Trypsin-EDTA (0.05%). DMEM containing 10% FBS was used to inactivate the trypsin. Cells and spent medium were centrifuged at 1000 rpm for three minutes. The cell pellet was washed by PBS thrice and resuspended in 1X Annexin V Binding Buffer. Cell suspension was transferred to a 5 mL culture tube or 96-well plate, where propidium iodide (PI) stain and Annexin V were added in according to the manufacturer’s instructions. BD Bioscience FacsAria III (San Jose, CA, USA) or Beckman Coulter Cytoflex S flow cytometer (Brea, CA, USA) was used to analyse the fluorescent output on the FITC and PI emission wavelengths. The fluorescence intensity of 10,000 cells was recorded. FlowJo 7.6 or CytoFLEX CytExpert 2.3 software was used for data analysis.

### Colony formation assay

Cells (10^6^ cells/well) were seeded to 6-well plates and treated with Sch B or 5-FU for 48 hours. At the end of treatment, treated cells were trypsinised and 500 cells were replated to clean 6-well plates and allowed to grow for an additional 14 days. Formed colonies were fixed with 100% ethanol, then stained with 1% crystal violet before being captured and counted manually.

### BrdU-ELISA assay

Cells were seeded at 10^4^ cells/well concentration onto 96-well culture plates and treated with Sch B for 48 hours. BrdU Cell Proliferation ELISA Kit (ab126556, abcam) was used according to manufacturer’s instructions. 10 μL of 10 × BrdU Solution was added to the cells, which was then incubated for four hours. After washed with PBS, the cells were fixed with fixing solution. Next, anti-BrdU monoclonal detector antibody (100 μL/well) was added to incubate the plate for one hour at room temperature. Finally, spectrophotometric microtiter plate reader measured the 450 nm absorbance value to identify the cell proliferation. BrdU incorporation of each sample was calculated as OD of the treatment sample minus the OD mean of control without addition of BrdU.

### Real-time quantitative polymerase chain reaction (RT-qPCR)

HCT-116 cells (6 x 10^5^ cells/well) were treated with various concentrations of Sch B for 48 hours. RNAiso Plus or GeneJET RNA Purification kit was used to extract total RNA, according to the manufacturer’s instructions. Purified RNA was resuspended in 50 μL of nuclease-free water and stored at −80 °C. RNA concentrations were measured using a NanoDrop ND-1000 Spectrophotometer (Nano-Drop Technologies, Wilmington, DE). RNA quality was determined by ensuring a value of 1.8 to 2.0 for the A260/A280 ratio. Complementary DNA (cDNA) was prepared from 500 ng of total RNA using the HiScript^TM^ RT SuperMix or PrimeScript RT reagent kit with gDNA Eraser for qPCR according to the manufacturer’s instructions. qPCR was performed to quantify the mRNA expression levels of ER stress-(*DDIT3, ATF3, ATF5, SMAD3*, HSPA5) and apoptosis-related genes (*BAX, BCL2, CASP3*). Glyceraldehyde-3-phosphate dehydrogenase (GAPDH) levels was also assessed to serve as an internal control for RNA integrity and loading. No differences in GAPDH amount between the groups were found for all the genes investigated (data not shown). All samples were run on a StepOnePlus Real-Time PCR system (Applied Biosystems, Foster City, CA) or LightCycler ® 96 System (Roche, Burgess Hill, UK) using 2 µl of cDNA and AceQ qPCR SYBR Green Master Mix or TB Green® Premix Ex Taq^TM^, with final primer concentrations of 0.5 µM per primer in a final volume of 10 µL. PCR was run using the default fast program (45 cycles of 95 °C for 5 s and 60 °C for 30 s). To ensure the reliability of RT-qPCR data, the amplicons are kept short (<250 bp). The amplification specificity was checked with melting curve analysis and gel electrophoresis. All PCR reactions were performed in duplicate. Relative changes in gene expression levels in cultured intestinal cells were analysed using the 2^-△△CT^ method as described previously [50]. Human-specific primers are described in Supplementary Table 1.

### Protein extraction, SDS-PAGE and Western Blot

Cells were seeded to 60mm culture dishes or 6-well plates and treated with Sch B for 48 hours. Cells were lysed with RIPA lysis buffer, supplemented with protease inhibitor cocktail, and total protein was extracted. Protein levels were normalized using DC Protein Assay. 20-50µg proteins were loaded to 10% SDS-PAGE gel and blotted to a PDVF membrane. The membrane was blocked in 5% dry milk (w/v) in Tris-buffered saline (TBS) containing 0.05% (v/v) Tween 20 (TBST) buffer for 1 hour at room temperature. Proteins were probed with specific primary antibodies overnight at 4°C. The membrane was washed with TBST for five times, five minutes each, and incubated with secondary antibodies for one hour at room temperature. The membrane was washed with TBST five times prior to visualization by Clarity Western ECL blotting kit (BioRad). Chemiluminescence was detected with a digital imaging system (ChemiDoc XRS+ system with image lab software, BioRad). The band intensity was quantified by ImageJ (NIH, Bethesda, Maryland, USA).

### Raman spectroscopy

The Sch B powder sample was first analysed with a conventional benchtop Renishaw InVia Qontor Raman spectrometer (Renishaw, Wotton-under-Edge, Gloucestershire, UK) with an excitation line of 785 nm and sample acquisition of 10 s. Cells were treated with or without Sch B and were lysed in ice-cold DMEM followed by passing through syringe needles at least three times. The samples were then measured by using an in-house compact Raman spectrometer with an excitation line of 785 nm. Although the in-house system had lower spectral resolution than the conventional one, the in-house Raman system provided the flexibility and ease of measuring cellular suspensions directly with higher reproducibility. The measurements used a quartz cuvette sample holder and a signal acquisition of 100 s.

### Computational docking

Molecular docking of the Sch B molecule into the CHOP protein was carried out using the Molecular Operating Environment (MOE) program. The Sch B molecule was built using the ‘Builder’ function in MOE and the structure for the CHOP protein was obtained from the AlphaFold Protein Structure Database (AlphaFold DB) (Uniprot ref: P35638). Docking sites were identified using the ‘SiteFinder’ tool, which identifies potential sites based on an Alpha spheres approach. Docking was performed using the ‘Dock’ module, using the Triangle Matcher placement methodology and 100 placement poses, with optimisation carried out using a molecular mechanics method using the Amber 10 forcefield. The binding energy (E_refine) was determined from the change in the non-bonded energy between ligand and receptor upon addition of the ligand. The binding free energy (E_score2) was determined using the GBVI/WSA scoring function.

### siRNA transfection

Cells were siRNA transfected, using Lipofectamine® RNAiMAX reagent, according to the manufacturer’s instructions. Knockdown efficiency was validated by qRT-PCR and Western blot analysis. After a 72-h incubation, transfected cells were treated without or with Sch B for 48 hours and then harvested for RNA and protein extraction, CCK cell viability assay and Annexin V/PI flow cytometry.

### Immunofluorescence

HCT116 cells were fixed (3.7% formaldehyde, 10 minutes), permeabilized (0.25% Triton X-100, 5% FBS, 15 minutes), blocked (5% FBS, 1 hour at RT), incubated with anti-DDIT3/GADD153/CHOP (ab56107, Abcam) (1:200) in 5% FBS overnight at 4°C and Alexa Fluor® 488 goat anti-mouse IgG secondary antibody (1:200; #10696113, ThermoFisher) for one hour at RT. Slides were counter-stained and mounted (Fluoromount-G™ Mounting Medium, #5596276, ThermoFisher), imaged by a Zeiss Axiovert LSM 710 VIS40S confocal microscope and quantified by ImageJ software.

### Mouse xenograft model of human colon cancer

Five-week-old BALB/c nude mice, weighing 18 to 20g, were obtained from the Laboratory Animal Unit. The animals were housed in individually ventilated cages. Temperature was kept at 22±2 °C, with 12 hours light/dark cycle. Animals were fed with standard diet (AIN-93G, Research Diets, USA) and given water *ad libitum*. After one week acclimatization, HCT-116 cells were injected subcutaneously to the mice. HCT116 cells were suspended in PBS at 1x10^7^ cells/ml concentration. 200μL of the cell suspension was injected to the right flank of the mice. One week was allowed for the tumour to form. Then, the mice were randomized into different groups, including PBS (Sham), Sch B (50mg/kg), 5-FU positive control (75mg/kg). Sch B was administered by oral gavage every other day, while 5-FU was given by intraperitoneal injection once a week (start of the week). The treatment lasted for 14 days, and body weight, tumour volume and faeces were observed throughout the period. All study protocols have been approved by the Committee on the Use of Live Animals in Teaching and Research (CULATR no. 5068-19) of the University of Hong Kong and the Department of Health of the HKSAR Government.

### Haematoxylin-Eosin staining

Tumours were harvested at the end of the sacrifice and fixed in 10% neutral buffered formalin at 4°C overnight. Fixed tumour tissues were then embedded in paraffin wax using Leica Tissue Processor (model ASP300S, Leica Microsystems Inc. Wetzlar, Germany). Haematoxylin and Eosin (H&E) staining was performed on 5-μm sections that were prepared and stained using kit (Vector Laboratories, CA, USA) according to manufacturer’s instructions. Stained slides were mounted with Histomount.

### RNA-seq

Total RNA was extracted using the Qiagen RNAeasy Kit (Qiagen, 74004). Quantity and quality were determined by Agilent 2100 bioanalyzer (Agilent Technologies, G2939A) before proceeding with RNA-seq library preparation. Paired-end sequencing with 100 bases read length was performed using BGI DNBSEQ™ platform (Wuhan, China). Sequence reads were mapped using HISAT2 using default settings and reads were aligned to human reference genome Hg37.2.

### Bioinformatic analysis

Log_2_ FC > 1.5 was used as cut-off for bioinformatic analysis. Differentially expressed genes, functional changes and activated canonical pathways were analysed with using Ingenuity Pathways Analysis (IPA, Qiagen).

### Statistical analysis

GraphPad Prism 9 was used for statistical analysis (GraphPad, USA). Normality tests were performed to determine whether parametric or non-parametric test was used for analyses. For parametric data, either two-tailed Student t-test, One-way or Two-way ANOVA with Holm-Šídák’s multiple comparisons test was used. For nonparametric data, either the two-tailed Mann–Whitney U-test or Kruskal–Wallis test with Dunn’s correction was used and paired data were analysed by Wilcoxon matched-pairs signed-rank test. Data were displayed as mean ± SD. * indicated a *P*-value of less than 0.05. ** indicated *P* <0.01. *** indicated *P* < 0.001. Quantification of confocal microscopy data was based on mean fluorescence intensity in 50□cells from independent experiments.

## Supplementary Figure Legends

**Supplementary Fig. 1. Supplementary data for Fig. 1.**

Effect of Sch B on viability of human colon cancer cell lines (HT29, SW620, SW480, LS174T, Caco-2) and a normal human intestinal cell line (CCD841 CoN), quantified by the Cell Counting Kit-8 (CCK-8) assay. Values were presented as mean ± SD, analysed by Kruskal–Wallis test with Dunn’s correction. \**P* < 0.05, \*\**P* < 0.01, \*\*\**P* < 0.001, compared with the control (i.e., 0 μM Sch B).

**Supplementary Fig. 2. Raman spectroscopy analysis of the uptake of Sch B and effect on human colon cancer cells. a,** The Raman spectrum of Sch B powder samples with several signature peaks occurred at 680, 714 and 1420 cm^-1^ (depicted with pink bars). **b,** The spectra of cell lysates and cell-free supernatants from cells cultured in the presence or absence of Sch B. Cell lysates and cell-free supernatants were obtained from human colon cancer cells exposed to 50 µM Sch B (i.e., 20 µg/mL) and analysed by an in-house compact Raman spectrometer. The degree of Sch B uptake to the cells after 0 and 48 hours were quantified from the relative intensities of the Raman peaks of Sch B (683, 719 and 1420 cm^-1^).

**Supplementary Fig. 3. Supplementary data for Fig. 6. a,** Top canonical pathways affected by Sch B in tumours of nude mice. **b,** IPA analysis showing lack of activation of unfolded protein response pathway in tumours of Sch B treated nude mice. Orange, up-regulated genes; blue, down-regulated genes.

**Supplementary Table 1.** qRT-PCR primers

**Supplementary Table 2.** Binding energies for Sch B at the nine docking sites identified in the CHOP protein using the site finder tool in MOE

## Contributions

Conceptualization – VC, HE, MW; Formal Analysis – VC, YL, BT, SJ, PD, PC, MW; Investigation – VC, HE, YL, BT, SJ, PD, PC, RA, MW; Methodology – VC, HE, PD, PC, RD, MW; Resources – HE, MW; Supervision – HE, MW; Funding acquisition – HE, MW; Visualization – VC, PD, PC, MW; Writing – original draft – VC, MW; Writing – review & editing – all authors.

## Corresponding authors

Correspondence to Hani El-Nezami or Murphy Lam Yim Wan

## Acknowledgements

The authors thank Drs. Steve Vayro and Robert Baldock at University of Portsmouth for technical support and advice on flow cytometry. The authors also thank Dr. Yihua Wang at University of Southampton for kindly providing the HCT116 cell line.

## Declarations

### Ethics approval and consent to participate

All study protocols have been approved by the Committee on the Use of Live Animals in Teaching and Research (CULATR no. 5068-19) of the University of Hong Kong and the Department of Health of the HKSAR Government.

### Consent for publication

All authors have read and approved of its submission to this journal.

### Availability of data and material

The data presented in this study are available on request from the corresponding author.

