## Supplementary Materials for "Schisandrin B suppresses colon cancer growth by inducing cell cycle arrest and apoptosis via the CHOP signalling pathway"

**Supplementary Fig. 1**

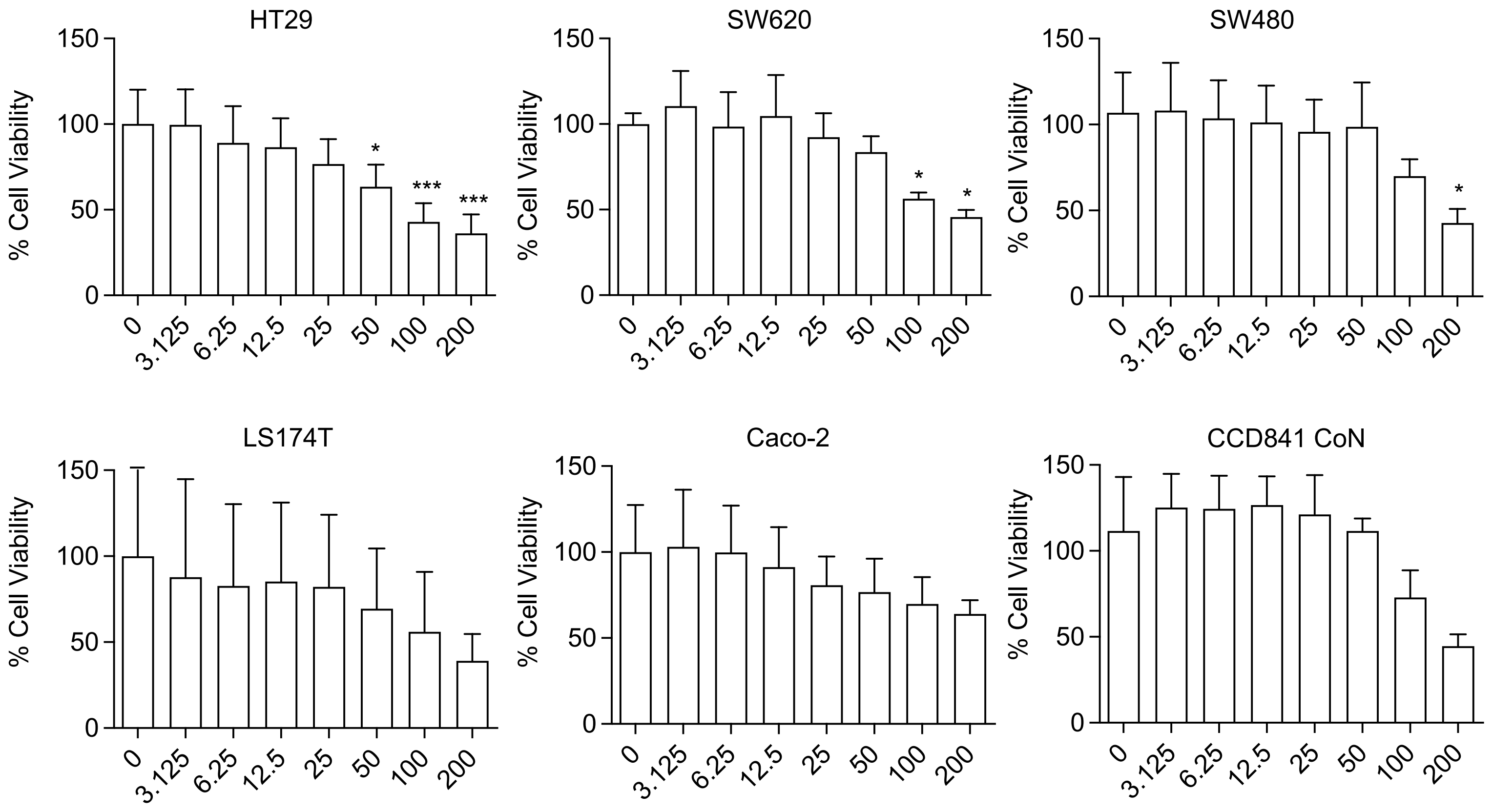

**Supplementary Fig. 2**

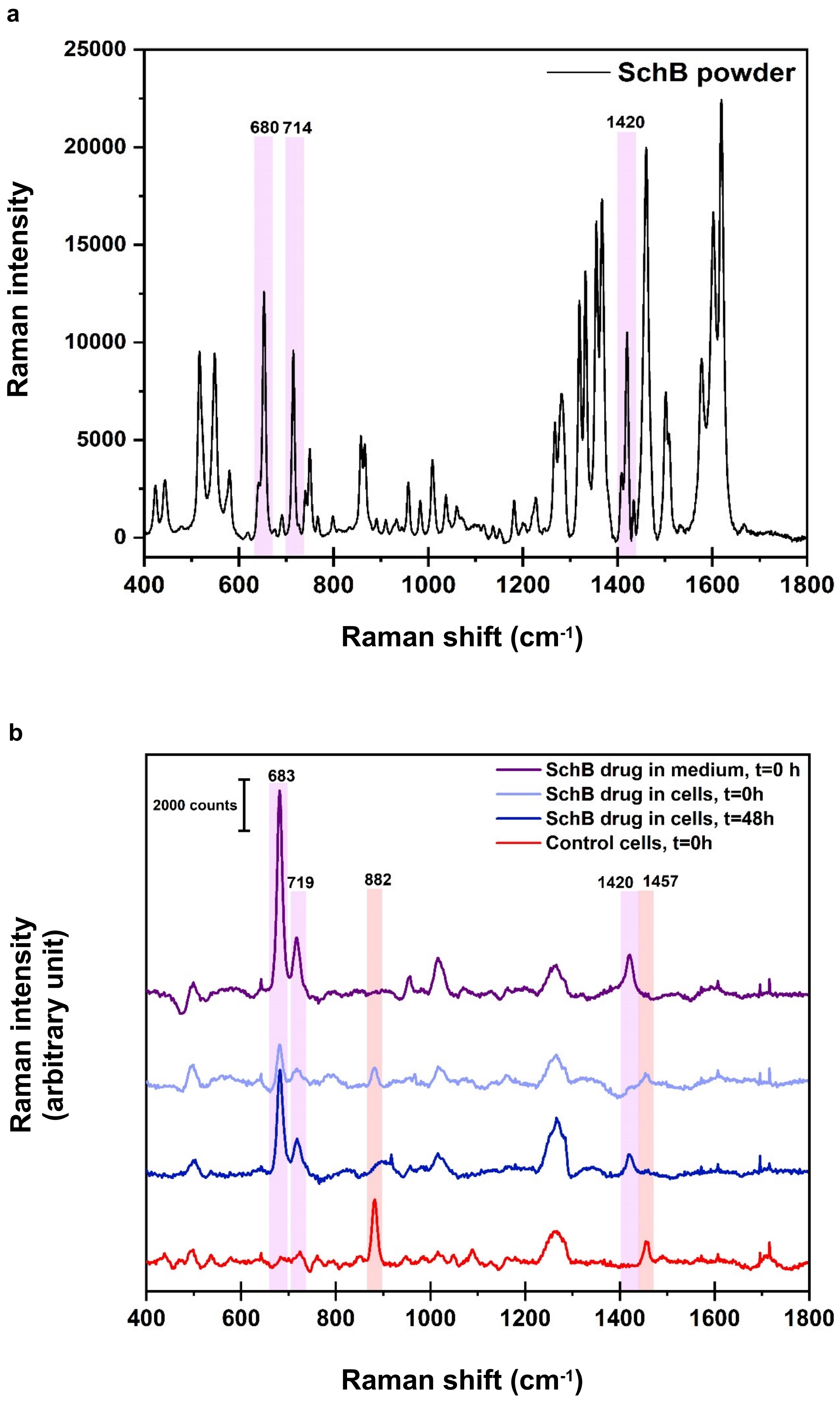

**Supplementary Fig. 3**

a

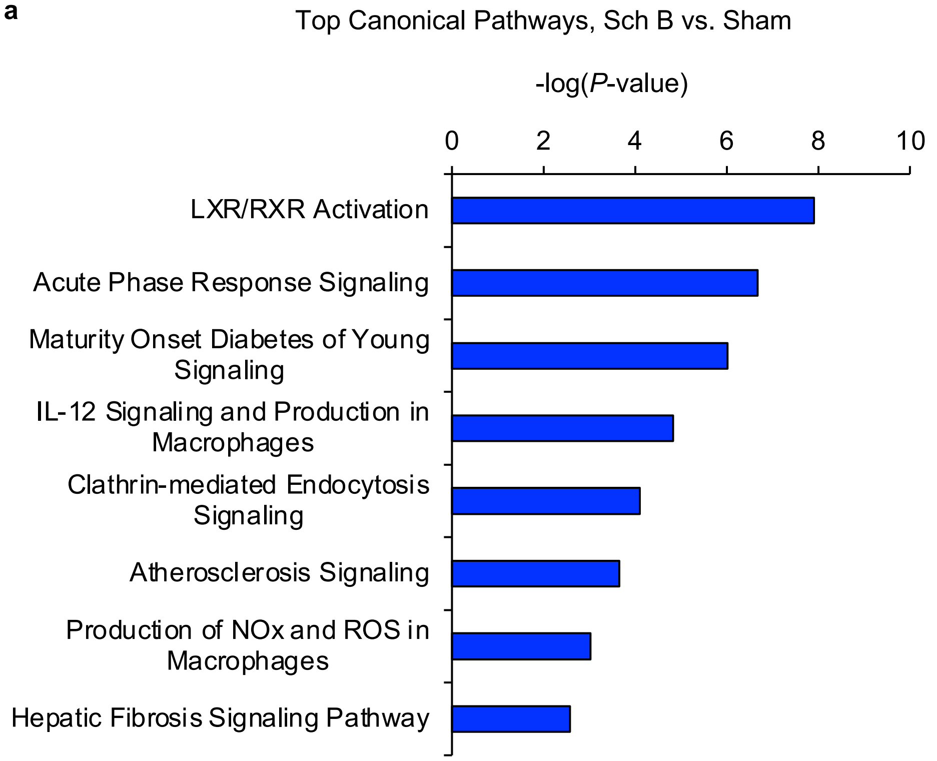


b

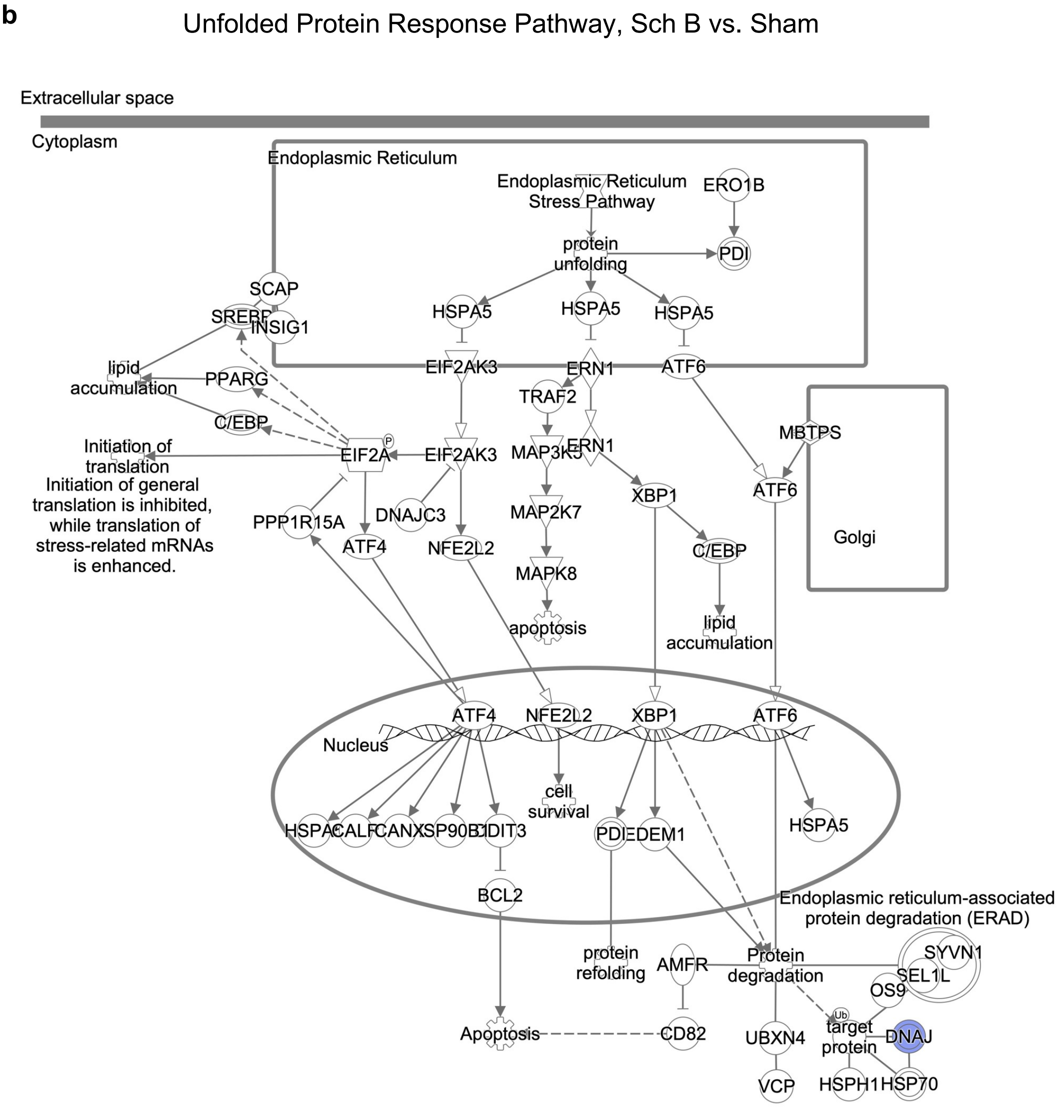

**Supplementary Table 1. qRT-PCR primers**

| **Gene** | **Primer sequences** | **Reference/Accession Number** |
|---|---|---|
| *BAX* | Fwd 5’-CCTGTGCACCAAGGTGCCGGAACT-3’<br>Rev 5’-CCACCCTGGTCTTGGATCCAGCCC-3’ | Elmaksoud *et al.* 2021, Biomedicine & Pharmacotherapy |
| *BCL2* | Fwd 5’-ATCGCCCTGTGGATGACTGAGT-3’<br>Rev 5’-GCCAGGAGAAATCAAACAGAGGC-3’ | NM_000633 |
| CASP3 | Fwd 5’-GGAAGCGAATCAATGGACTCTGG-3’<br>Rev 5’-GCATCGACATCTGTACCAGACC-3’ | NM_004346 |
| *DDIT3* | Fwd 5’-GGTATGAGGACCTGCAAGAGGT-3’<br>Rev 5’-CTTGTGACCTCTGCTGGTTCTG-3’ | NM_004083 |
| *SMAD3* | Fwd 5’-TGAGGCTGTCTACCAGTTGACC-3’<br>Rev 5’-GTGAGGACCTTGTCAAGCCACT-3’ | NM_005902 |
| *ATF3* | Fwd 5’-CGCTGGAATCAGTCACTGTCAG-3’<br>Rev 5’-CTTGTTTCGGCACTTTGCAGCTG-3’ | NM_001674 |
| *ATF5* | Fwd 5’-GCTCGTAGACTATGGGAAACTCC-3’<br>Rev 5’-CATCCAGTCAGAGAAGCCATCAC-3’ | NM_012068 |
| *HSPA5* | Fwd 5’-CTGTCCAGGCTGGTGTGCTCT-3’<br>Rev 5’-CTTGGTAGGCACCACTGTGTTC-3’ | NM_005347 |
| *GAPDH* | Fwd 5’-ACCAGCCCCAGCAAGAGCACAAG-3’<br>Rev 5’-TTCAAGGGGTCTACATGGCAACTG-3’ | Wang *et al.* 2006, Journal of Immunology |

**Supplementary Table 2. Binding energies for Sch B at the nine docking sites identified in the CHOP protein using the site finder tool in MOE**

| Site Number | Binding Energy kcal/mol | Binding Free Energy kcal/mol | Residues |
|---|---|---|---|
| 1 | -22.90 | -5.07 | PRO7 PHE8 SER9 PHE10 GLY11 THR12 LEU13 |
| 2 | -23.63 | -4.97 | ARG140 GLN143 GLU144 GLU146 ARG147 ARG150 |
| 3 | -28.54 | -5.39 | ASN44 GLU45 GLU46 GLU47 GLU48 SER49 |
| 4 | -22.97 | -5.01 | MET120 LYS121 LYS123 GLU124 ASN127 |
| 5 | -20.43 | -4.73 | VAL28 LEU29 ASP32 GLU33 |
| 6 | -20.71 | -4.71 | GLN133 GLU136 GLU137 ARG140 |
| 7 | -27.49 | -5.31 | GLU46 GLU47 SER49 LYS50 |
| 8 | -21.55 | -4.74 | ALA115 GLN118 ARG119 |
| 9 | -20.80 | -4.80 | LEU13 GLU17 LEU18 TRP21 |
